# Risk of preeclampsia in patients with genetic predisposition to common medical conditions: a case-control study

**DOI:** 10.1101/2020.03.04.976472

**Authors:** Kathryn J. Gray, Vesela P. Kovacheva, Hooman Mirzakhani, Andrew C. Bjonnes, Berta Almoguera, Melissa L. Wilson, Sue Ann Ingles, Charles J. Lockwood, Hakon Hakonarson, Thomas F. McElrath, Jeffrey C. Murray, Errol R. Norwitz, S. Ananth Karumanchi, Brian T. Bateman, Brendan J. Keating, Richa Saxena

**Author notes:** Department of Medicine, Channing Division of Network Medicine, Brigham and Women’s Hospital, Boston, MA. Department of Biomedical Informatics, Harvard Medical School, Boston, MA. Department of Anesthesiology, Brigham and Women’s Hospital, Boston, MA. **CORRESPONDING AUTHOR:** Kathryn J. Gray, MD PhD, Division of Maternal-Fetal Medicine, Brigham and Women’s Hospital, 75 Francis St, CWN-3, Boston, MA, 02115, USA.

## Abstract

**Objective:** To assess whether women with a genetic predisposition to medical conditions known to increase preeclampsia risk have an increased risk of preeclampsia in pregnancy.

**Design:** Case-control study.

**Setting and population:** Preeclampsia cases (n=498) and controls (n=1864) of European ancestry from 5 US sites genotyped on a cardiovascular gene-centric array.

**Methods:** Significant single nucleotide polymorphisms (SNPs) from 21 traits in 7 disease categories (cardiovascular, inflammatory/autoimmune, insulin resistance, liver, obesity, renal, thrombophilia) with published genome-wide association studies (GWAS) were used to create a genetic instrument for each trait. Multivariable logistic regression was used to test the association of each continuous, scaled genetic instrument with preeclampsia. Odds of preeclampsia were compared across quartiles of the genetic instrument and evaluated for significance using a test for trend.

**Main Outcome Measures:** preeclampsia.

**Results:** An increasing burden of risk alleles for elevated diastolic blood pressure (DBP) and increased body mass index (BMI) were associated with an increased risk of preeclampsia (DBP: overall OR 1.11 (1.01-1.21), p=0.025; BMI: OR 1.10 (1.00-1.20), p=0.042), while risk alleles associated with elevated alkaline phosphatase (ALP) were protective (OR 0.89 (0.82-0.97), p=0.008), driven primarily by pleiotropic effects of variants in the *FADS* gene region. The effect of DBP genetic loci was even greater in early-onset (<34 weeks) preeclampsia cases (OR 1.30 (1.08-1.56), p=0.005). For all other traits, the genetic instrument was not robustly associated with preeclampsia risk.

**Conclusions:** These results suggest that the underlying genetic architecture of preeclampsia is shared with other disorders, specifically hypertension and obesity.

**TWEETABLE ABSTRACT:** Genetic predisposition to increased diastolic blood pressure and obesity increases the risk of preeclampsia.

## INTRODUCTION

Preeclampsia (PE) is a severe, pregnancy-specific disorder affecting 3-8% of all gestations and characterized by new-onset hypertension and proteinuria after 20 weeks gestation. Despite extensive investigation, the etiology remains poorly understood and PE continues to be a leading cause of maternal and neonatal morbidity and mortality.

Genetics influence PE risk—multiple epidemiologic studies estimate the heritability at 55-60%, of which 30-35% is maternal and 20% fetal^1–4^, with early-onset PE having the largest genetic component and environmental factors contributing more to late-onset PE^5^. Understanding PE heritability is challenging due to the involvement of two genomes (maternal and fetal), the heterogeneous nature of the disease, and limited availability of large, well-phenotyped PE cohorts^6^. Recently, the first fetal and maternal PE genome-wide association studies (GWAS) have been published revealing a few genetic loci contributing to risk^7–11^; however, the majority of PE heritability remains unexplained.

Co-existing maternal medical conditions (e.g., diabetes, chronic hypertension, renal disease, autoimmune disease, antiphospholipid antibody syndrome) also increase PE risk^12,13^. Many additional clinical risk factors for PE are well-established, including obesity, dyslipidemia, multifetal gestation, nulliparity, use of assisted reproductive technology, previous PE, and family history of PE or cardiovascular disease^12–17^. Specifically, women with chronic hypertension have a 16-25% risk of PE^12,13^, and women with prior PE are more likely to use antihypertensive medications both the short-and long-term^18^. For obesity, the higher the BMI, the greater the risk of PE; women who are obese (BMI >35) have a 3-fold increased risk^19–23^. For dyslipidemia, PE is associated with elevated total cholesterol, non-HDL cholesterol, and triglycerides throughout pregnancy, as well as with lower HDL levels in the third trimester^16,24,25^. Additionally, following a pregnancy with PE, women have a 2-4-fold increased lifetime risk of cardiovascular disease (CVD)^26–29^.

Given this, we hypothesized that women with genetic predisposition to these medical conditions would have an increased risk of PE. To test this hypothesis, we identified single nucleotide polymorphisms (SNPs) for relevant traits from the largest published European genome-wide association studies (GWAS) for each trait and tested whether these SNPs increased PE risk in a maternal case/control sample of European ancestry from the United States genotyped using the ITMAT-Broad-CARe (IBC) genotyping array, that captures genetic diversity across >2000 candidate gene regions related to cardiovascular, inflammatory, and metabolic phenotypes^30^.

## METHODS

### Study population

We utilized maternal GWAS data from a previously assembled IRB-approved cohort^8^ with European-ancestry cases (n=498) and controls (n=449) from five US cities and population controls from the National Heart, Lung, and Blood Institute (NHLBI) studies, Atherosclerosis Risk in Communities (ARIC; n=645)^31^ and Coronary Artery Risk Development in Young Adults (CARDIA; n=770)^32^, for a total of 498 cases and 1864 controls (Table 1). The published GWAS describes the specifics of the maternal sample collection, genotyping, and quality control for this cohort^8^. PE status at all sites was defined by standard American Congress of Obstetrician and Gynecologists (ACOG) criteria^33^. Controls from the US sites were normotensive pregnant women without medical co-morbidities.

**Table 1.**
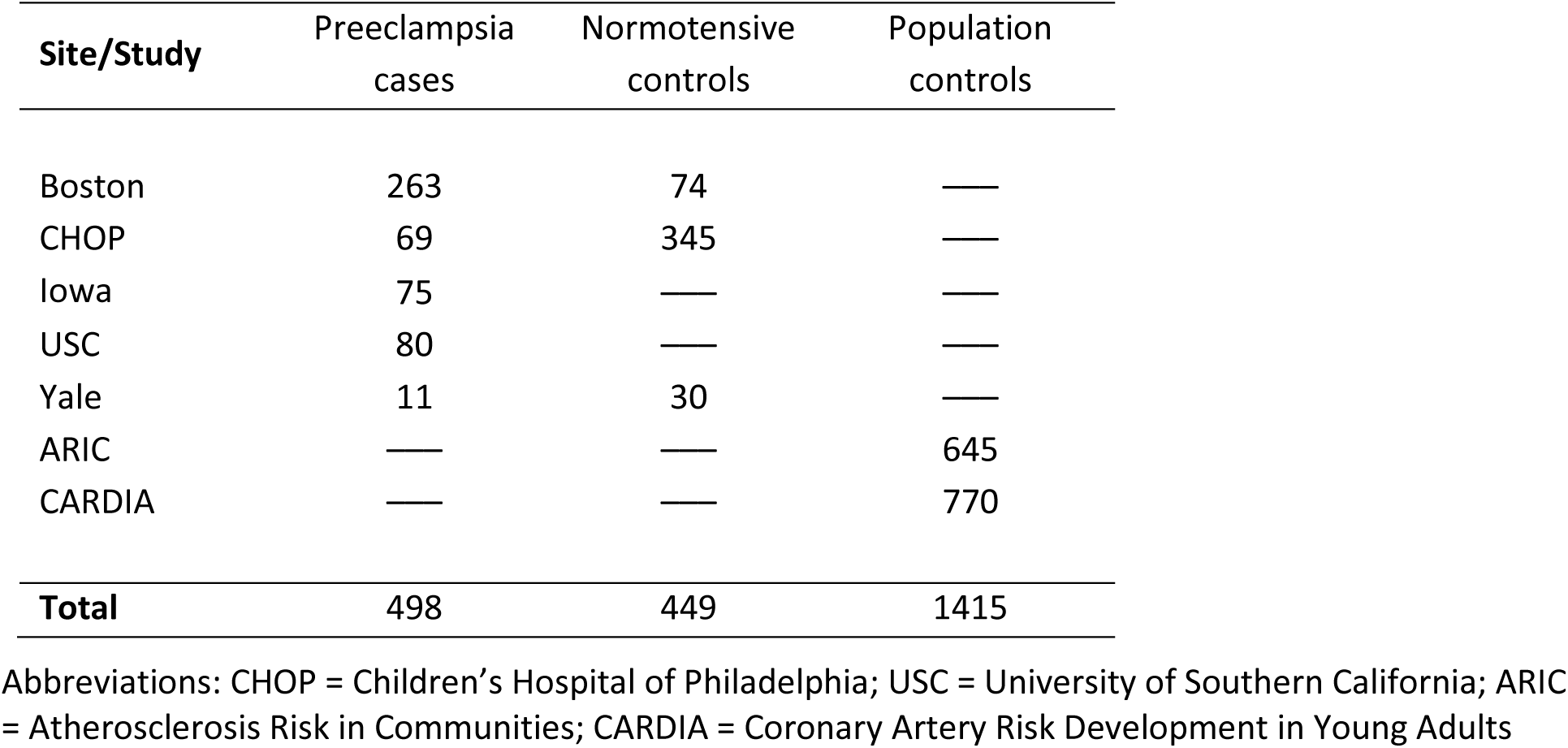
Collection site of European ancestry preeclampsia cases (n=498) and controls (n=1864).

| Site/Study | Preeclampsia cases | Normotensive controls | Population controls |
| --- | --- | --- | --- |
| Boston | 263 | 74 | — |
| CHOP | 69 | 345 | — |
| Iowa | 75 | — | — |
| USC | 80 | — | — |
| Yale | 11 | 30 | — |
| ARIC | — | — | 645 |
| CARDIA | — | — | 770 |
| **Total** | **498** | **449** | **1415** |

Abbreviations: CHOP = Children's Hospital of Philadelphia; USC = University of Southern California; ARIC = Atherosclerosis Risk in Communities; CARDIA = Coronary Artery Risk Development in Young Adults

### Identification of traits and their associated single nucleotide polymorphisms (SNPs)

The GWAS Catalog, curated by the National Human Genome Research Institute (NHGRI) and the European Bioinformatics Institute (EBI), was used to identify genome-wide association studies performed up to the year 2017 in individuals of European ancestry for traits known to be associated with PE^34^. We searched specifically for studies on traits in 7 categories: cardiovascular, inflammatory/autoimmune, insulin resistance, liver, obesity, renal, and thrombophilia (Table 2).

**Table 2.** Disease categories and specific traits assessed using risk SNPs.

| Disease category | Trait |
| --- | --- |
| Cardiovascular | Systolic blood pressure (SBP), diastolic blood pressure (DBP), total cholesterol, low-density lipoprotein (LDL), high-density lipoprotein (HDL), triglycerides, coronary artery disease (CAD), ischemic stroke |
| Inflammatory/ autoimmune | C-reactive protein (CRP), systemic lupus erythematosus (SLE) |
| Insulin resistance | Adiponectin, fasting blood glucose, type 2 diabetes mellitus (T2DM) |
| Liver | Alkaline phosphatase (ALP), gamma-glutamyl transferase (GTT) |
| Obesity | Body mass index (BMI), birthweight |
| Renal | Estimated glomerular filtration rate from serum creatinine (eGFR<sub>Cr</sub>), uric acid |
| Thrombophilia | Fibrinogen, venous thromboembolism (VTE) |

Within the cardiovascular category, we identified GWAS for systolic and diastolic blood pressure^35^, total cholesterol^36,37^, low-density lipoprotein (LDL)^36,37^, high-density lipoprotein (HDL)^36,37^, triglycerides^36,37^, coronary artery disease (CAD)^38,39^, and ischemic stroke^40^. PE is highly associated with hypertension both prior to and following pregnancy^12,13^. Other cardiovascular risk factors including increased levels of total cholesterol, LDL-C, and triglycerides and decreased levels of HDL-C have also been associated with PE^24,41^. The increased long-term risk of cardiovascular disease in women with prior PE includes an increased risk of CAD, ischemic stroke, heart failure, cardiac procedures, and cardiovascular-related hospitalizations^26–29,42,43^.

Within the inflammatory/autoimmune category, we identified GWAS for C-reactive protein (CRP)^44^ and systemic lupus erythematosus (SLE)^45^. Women with increased CRP are at significantly increased risk of PE, even after adjustment for clinical confounders such as BMI^46^. Women with SLE are at increased risk of PE, particularly early-onset PE, even after adjustment for chronic hypertension and antiphospholipid antibody syndrome (APS)^47^. While lupus nephritis, APS, and chronic hypertension with SLE are associated with the highest risk of PE, even uncomplicated SLE in remission is associated with increased risk^48^.

Within the insulin resistance category, we identified GWAS for adiponectin^49^, fasting glucose^49^, fasting insulin^50^, type 2 diabetes mellitus (T2DM)^51^, and polycystic ovarian syndrome (PCOS)^52^. Women with pregestational or gestational diabetes have a 2-4 fold increased PE risk and non-diabetic women with PE are at increased risk of later T2DM^53^. Adiponectin levels are associated with the development of gestational diabetes^54,55^. Women with PCOS have a high rate of metabolic syndrome and significantly higher risks of adverse pregnancy outcomes including PE^56^.

Within the liver category, we identified GWAS for alkaline phosphatase (ALP)^57^, gamma-glutamyl transferase (GTT)^57^, and alanine aminotransferase (ALT)^57^. Women with liver dysfunction during pregnancy (e.g., intrahepatic cholestasis of pregnancy) are at increased risk for PE^58^. Also, transaminitis is a key diagnostic feature for particular PE subtypes, including PE with severe features and HELLP (hemolysis, elevated liver enzymes, low platelets) syndrome^33^.

Within the thrombophilia category, we identified GWAS for fibrinogen^59^ and venous thromboembolism (VTE)^60^. PE is associated with a state of hypercoagulability. Specifically, fibrinogen levels are elevated in women with PE^61^ and women with PE are at significantly increased risk of VTE^42,62^. Although antiphospholipid antibody syndrome is a strong clinical risk factor for PE, the only published GWAS is a small study of Japanese women^63^.

Within the obesity category, we identified GWAS for body mass index (BMI)^64^ and maternal birthweight^65^. Increased BMI is a prevalent risk factor for PE and the degree of obesity is directly correlated to the level of risk^19–23^. We included maternal birthweight as women who are small at birth and obese as adults have a particularly high risk of PE^66^.

Within the renal category, we identified GWAS for estimated glomerular filtration rate from serum creatinine (eGFRCr)^67^ and uric acid^68^. Women with decreased GFR prior to pregnancy are at increased risk of PE^69,70^, and women with PE have a reduced GFR^71,72^. In addition to its role as a marker for renal function, elevations in uric acid are associated with PE and may contribute directly to disease pathogenesis^73–75^.

For each of the above traits, the most recent and largest published GWAS (up to the year 2017) for individuals of European ancestry was identified and trait-associated SNPs with *p*-values ≤ 1.0 x 10^−6^ were curated (see Supplementary Tables S1-S21 for each trait). For each curated SNP, we determined if the SNP was genotyped on the IBC array used in our cohort^30^. If the SNP was not present on the array, we determined if a proxy SNP was present in the 1000 Genomes phase 3 (1KGPv3) CEU population^76^ using SNAP, the SNP Annotation and Proxy Search available through the Broad Institute^77^. For each proxy, the proxy SNP with the strongest correlation was chosen. Most proxy SNPs had r^2^>0.8, and only SNPs with proxies of r^2^>0.5 were included in our analyses (see Supplementary Tables S1-S21). Using this approach, the noted number of SNPs were available for each trait (Table 3).

**Table 3.** Preeclampsia risk by quartiles of trait-specific risk SNPs.

| TRAIT* | Q1<br>odds ratio<br>(Ref) | Q2<br>odds ratio<br>(95% CI) | Q3<br>odds ratio<br>(95% CI) | Q4<br>odds ratio<br>(95% CI) | Overall<br>odds ratio<sup>†</sup><br>(95% CI) | p-value<br>(for trend<br>by quartile) |
| --- | --- | --- | --- | --- | --- | --- |
| **Cardiovascular** |  |  |  |  |  |  |
| **SBP**<br>(19 SNPs) | 1.00 | 0.82<br>(0.62-1.08) | 0.88<br>(0.67-1.15) | 0.84<br>(0.64-1.10) | 0.96<br>(0.88-1.04) | 0.322 |
| **DBP**<br>(16 SNPs) | 1.00 | 1.37<br>(1.03-1.82) | 1.42<br>(1.07-1.89) | 1.39<br>(1.05-1.86) | 1.11<br>(1.01-1.21) | **0.025** |
| **Total cholesterol**<br>(20 SNPs) | 1.00 | 1.03<br>(0.78-1.36) | 0.97<br>(0.73-1.28) | 1.10<br>(0.84-1.45) | 1.02<br>(0.94-1.12) | 0.589 |
| **LDL**<br>(12 SNPs) | 1.00 | 0.95<br>(0.71-1.26) | 1.20<br>(0.91-1.58) | 1.13<br>(0.85-1.49) | 1.06<br>(0.98-1.16) | 0.164 |
| **HDL**<br>(28 SNPs) | 1.00 | 1.01<br>(0.77-1.32) | 0.70<br>(0.53-0.93) | 0.87<br>(0.66-1.14) | 0.93<br>(0.85-1.01) | 0.096 |
| **Triglycerides**<br>(15 SNPs) | 1.00 | 0.92<br>(0.70-1.21) | 0.94<br>(0.71-1.24) | 0.95<br>(0.72-1.26) | 0.98<br>(0.90-1.07) | 0.626 |
| **CAD**<br>(10 SNPs) | 1.00 | 1.18<br>(0.88-1.57) | 1.43<br>(1.08-1.90) | 1.11<br>(0.82-1.51) | 1.05<br>(0.96-1.15) | 0.294 |
| **Ischemic stroke**<br>(5 SNPs) | 1.00 | 0.95<br>(0.71-1.26) | 0.81<br>(0.60-1.10) | 0.97<br>(0.72-1.31) | 0.97<br>(0.89-1.07) | 0.584 |
| **Inflammatory/ autoimmune** |  |  |  |  |  |  |
| **CRP**<br>(9 SNPs) | 1.00 | 0.93<br>(0.70-1.23) | 1.10<br>(0.84-1.44) | 0.97<br>(0.73-1.28) | 1.00<br>(0.92-1.10) | 0.949 |
| **SLE**<br>(9 SNPs) | 1.00 | 1.12<br>(0.85-1.48) | 1.04<br>(0.78-1.38) | 1.09<br>(0.82-1.44) | 1.01<br>(0.93-1.11) | 0.775 |
| **Insulin resistance** |  |  |  |  |  |  |
| **Adiponectin**<br>(3 SNPs) | 1.00 | 1.06<br>(0.74-1.51) | 0.95<br>(0.67-1.35) | 0.95<br>(0.66-1.36) | 0.97<br>(0.87-1.07) | 0.521 |
| **Fasting glucose**<br>(5 SNPs) | 1.00 | 0.75<br>(0.56-1.00) | 0.92<br>(0.69-1.23) | 0.86<br>(0.65-1.15) | 0.99<br>(0.90-1.08) | 0.782 |
| **T2DM**<br>(21 SNPs) | 1.00 | 1.17<br>(0.89-1.54) | 0.94<br>(0.71-1.24) | 0.97<br>(0.73-1.29) | 0.97<br>(0.89-1.06) | 0.470 |
| **Liver** |  |  |  |  |  |  |
| **ALP**<br>(5 SNPs) | 1.00 | 0.85<br>(0.65-1.11) | 0.63<br>(0.47-0.84) | 0.74<br>(0.56-0.97) | 0.89<br>(0.82-0.97) | **0.008** |
| **GTT**<br>(6 SNPs) | 1.00 | 1.29<br>(0.97-1.72) | 1.31<br>(0.99-1.74) | 1.19<br>(0.90-1.57) | 1.05<br>(0.97-1.15) | 0.250 |
| **Obesity** |  |  |  |  |  |  |
| **BMI**<br>(15 SNPs) | 1.00 | 0.96<br>(0.72-1.27) | 1.14<br>(0.86-1.51) | 1.28<br>(0.97-1.69) | 1.10<br>(1.00-1.20) | **0.042** |
| **Birthweight**<br>(6 SNPs) | 1.00 | 1.07<br>(0.80-1.42) | 1.07<br>(0.80-1.44) | 1.17<br>(0.87-1.57) | 1.04<br>(0.95-1.13) | 0.457 |
| ***Renal*** |  |  |  |  |  |  |
| **eGFR<sub>Cr</sub>**<br>(9 SNPs) | 1.00 | 0.75<br>(0.57-0.99) | 0.86<br>(0.66-1.13) | 0.83<br>(0.63-1.09) | 0.96<br>(0.88-1.05) | 0.391 |
| **Uric acid**<br>(11 SNPs) | 1.00 | 1.24<br>(0.94-1.65) | 1.17<br>(0.88-1.55) | 1.16<br>(0.88-1.54) | 1.04<br>(0.95-1.14) | 0.379 |
| ***Thrombophilia*** |  |  |  |  |  |  |
| **Fibrinogen**<br>(5 SNPs) | 1.00 | 0.91<br>(0.68-1.21) | 0.94<br>(0.72-1.24) | 1.06<br>(0.81-1.40) | 1.03<br>(0.94-1.12) | 0.565 |
| **VTE**<br>(5 SNPs) | 1.00 | 0.99<br>(0.75-1.31) | 0.66<br>(0.49-0.89) | 0.89<br>(0.66-1.18) | 0.93<br>(0.85-1.02) | 0.101 |

\*Specific SNPs assessed for each trait are listed in the Online Supplement
† Odds ratio reflects the per risk allele increase in preeclampsia risk, adjusted for study site and principle components of ancestry

After curation of these GWAS and their associated SNPs, we had a total of 21 distinct traits within 7 categories with SNPs or proxy SNPs available (Table 2). Some traits of interest (alanine aminotransferase (ALT), fasting insulin, polycystic ovarian syndrome (PCOS)) could not be included in analysis as the implicated SNPs were not genotyped in our cohort. Of note, while newer GWAS have been published since 2017 for some of the selected traits, the additional loci reported could not able to be examined as our cohort was genotyped on the gene-centric IBC array (which has limited SNPs compared to newer arrays) and the additional loci or proxies were not available in our data.

### Statistical Analysis

#### Determination of genetic instruments

For each trait, we derived a genetic instrument to assess risk for each individual participant by summing the number of risk SNPs, which were weighted by the respective allelic effect size (β-coefficient) from the original discovery cohort using PLINK, as previously described^78–81^. If genotype data were missing for a particular individual for a particular SNP, then the expected value was imputed based on the sample allele frequency of the SNP (in total, this affected very few loci (<5%), as SNPs were removed for call rates <95%). Scaling of the genetic instrument for each trait was performed to allow interpretation of the effects on PE as a per-1 risk allele increase in the total sum of risk SNPs for each trait (division by twice the sum of the β-coefficients and multiplication by twice the SNP count representing the maximum number of risk alleles). Multivariable logistic regression in R was used to test the association of each continuous, scaled genetic instrument with PE, adjusted for study site and principal components of ancestry, leading to an adjusted, overall odds ratio. The odds of PE were compared across quartiles of each genetic instrument and evaluated for significance using a test for trend.

## RESULTS

The distribution of European ancestry PE cases and normotensive controls between the 5 US sites, as well as the population controls, is shown in Table 1. As noted previously, the cases in this sample are enriched for early-onset PE (diagnosis <37 weeks gestation; 40%), hemolysis-elevated liver enzymes-low platelets (HELLP) syndrome (29%), and PE with severe features (43%)^8^.

For each of the 21 traits identified, the derived genetic instrument was used to assess the risk of PE conferred by the trait-associated risk SNPs (Table 3). For the cardiovascular traits, we found that an increasing burden of risk alleles for elevated diastolic blood pressure (DBP) was associated with increased PE risk at all quartiles of risk with an overall OR of 1.11 (1.01-1.21), p=0.025, per risk allele. For coronary artery disease (CAD), patients in the third quartile of risk had a significantly increased risk of PE compared to the reference group (OR 1.43 (1.08-1.90), p=0.014), while for HDL, patients in the third quartile had a significantly decreased risk of PE compared to the reference (OR 0.70 (0.53-0.93), p = 0.014). In this analysis, we did not find an association of the genetic instruments for systolic blood pressure (SBP), total cholesterol, LDL, triglycerides, or ischemic stroke with PE. For the inflammatory/autoimmune traits, we did not find an association between CRP or SLE-associated genetic instruments and PE. Similarly, for the insulin resistance traits, adiponectin, fasting glucose, and type 2 diabetes (T2DM), the genetic instruments were not significantly associated with PE.

For the liver traits, gamma-glutamyl transferase (GTT)-associated risk alleles were not significantly associated with PE. However, an increasing burden of risk alleles for elevated alkaline phosphatase (ALP) levels was surprisingly protective for PE (overall OR 0.89 (0.82-0.97), p=0.008). As this result was not anticipated, we examined the particular SNPs (n=5) used to create the ALP genetic instrument in further detail (Supplementary Table S2; SNPs from most comprehensive GWAS of plasma liver enzyme concentrations^57^). Of these 5 SNPs, 2 SNPs contributed most significantly to the protective nature of the ALP variants: rs174601 (associated with *FADS1, FADS2*, and *C11orf10* gene expression) and rs579459 (near the *ABO* gene) (data not shown).

For the obesity traits, an increasing burden of risk alleles for increased body mass index (BMI) was associated with increased PE risk (overall OR 1.10 (1.00-1.20), p=0.042). The genetic instrument for maternal birthweight, however, was not associated with PE risk. For the renal traits, patients in the second quartile of risk for increased GFR, had a decreased risk of PE (OR 0.75 (0.57-0.99), p=0.044). In contrast, the genetic instrument for uric acid was not significantly associated with PE. For the thrombophilia traits, patients in the second quartile of genetic risk for VTE had a decreased risk of PE (OR 0.66 (0.49-0.89), p=0.006), while the genetic instrument for fibrinogen was not significantly associated with PE.

As early-onset PE has the strongest genetic predisposition^5^, we performed a sensitivity analysis for all 21 traits on the subset of PE cases with delivery < 34 weeks (n=103 cases) and < 37 weeks (n=201 cases). In the subset of cases with early-onset PE, the effect of DBP-associated risk alleles was greater than in the overall cohort (<34 weeks: overall OR 1.30 (1.08-1.56), p=0.005; <37 weeks: overall OR 1.19 (1.05-1.36), p=0.009) (Table 4). In contrast, the effects of BMI and ALP-associated risk alleles were diminished (BMI, <34 weeks: overall OR 1.04 (0.87-1.24), p=0.683; <37 weeks: overall OR 1.06 (0.93-1.21), p=0.400. ALP, <34 weeks: overall OR 0.86 (0.72-1.03), p=0.108; <37 weeks: overall OR 0.82 (0.72-0.94), p=0.004) (Table 4). For all other traits examined, the effects of the genetic instruments were not significant in the early-onset subgroup (data not shown).

**Table 4.** Early-onset preeclampsia risk by quartiles of trait-specific risk SNPs.

| **TRAIT\*** | **Q1**<br>**odds ratio**<br>**(Ref)** | **Q2**<br>**odds ratio**<br>**(95% CI)** | **Q3**<br>**odds ratio**<br>**(95% CI)** | **Q4**<br>**odds ratio**<br>**(95% CI)** | **Overall**<br>**odds ratio<sup>†</sup>**<br>**(95% CI)** | **p-value**<br>**(for trend**<br>**by quartile)** |
| --- | --- | --- | --- | --- | --- | --- |
| ***Delivery < 34 weeks*** |  |  |  |  |  |  |
| **DBP**<br>(16 SNPs) | 1.00 | 1.71<br>(0.89-3.28) | 1.91<br>(1.00-3.64) | 2.45<br>(1.31-4.58) | 1.30<br>(1.08-1.56) | **0.005** |
| **BMI**<br>(15 SNPs) | 1.00 | 0.76<br>(0.42-1.36) | 0.87<br>(0.50-1.51) | 1.03<br>(0.60-1.78) | 1.04<br>(0.87-1.24) | 0.683 |
| **ALP**<br>(5 SNPs) | 1.00 | 1.09<br>(0.65-1.84) | 0.66<br>(0.37-1.18) | 0.71<br>(0.40-1.27) | 0.86<br>(0.72-1.03) | 0.108 |
| ***Delivery < 37 weeks*** |  |  |  |  |  |  |
| **DBP**<br>(16 SNPs) | 1.00 | 1.73<br>(1.10-2.73) | 1.70<br>(1.08-2.68) | 1.91<br>(1.22-2.98) | 1.19<br>(1.05-1.36) | **0.009** |
| **BMI**<br>(15 SNPs) | 1.00 | 1.00<br>(0.65-1.52) | 0.97<br>(0.64-1.48) | 1.17<br>(0.78-1.77) | 1.06<br>(0.93-1.21) | 0.400 |
| **ALP**<br>(5 SNPs) | 1.00 | 0.81<br>(0.55-1.20) | 0.55<br>(0.36-0.85) | 0.60<br>(0.40-0.90) | 0.82<br>(0.72-0.94) | **0.004** |

\*Specific SNPs assessed for each trait are listed in the Online Supplement
<sup>†</sup> Odds ratio reflects the per risk allele increase in preeclampsia risk, adjusted for study site and principle components of ancestry

## DISCUSSION

### Main Findings

Here we detail the first comprehensive analysis of the effect of genetic predisposition for 21 clinical traits on PE risk, utilizing data from the largest published US maternal PE GWAS^8^. Among the 21 traits examined, risk alleles for elevated DBP and increased BMI were most robustly associated with PE risk, while risk alleles associated with elevated ALP levels were protective. Additionally, we identified a suggestive risk effect for coronary artery disease and possible protective effects for increased HDL, GFR, and VTE risk. The effect of DBP-associated risk alleles was strongest in early-onset PE cases, while the effects of BMI and ALP-associated risk alleles were diminished in this subset, suggesting that these effects are more important in late-onset disease. For the other traits examined, we did not find significant associations.

### Strengths and Limitations

Strengths of this study include utilization of the largest US-based maternal PE cohort with genome-wide SNP data reported to date and a comprehensive examination of PE-associated traits. This cohort is enriched with early-onset and severe cases that are more likely to have a genetic predisposition to PE^5^.

Despite its size relative to many other PE genetic studies, this cohort has a limited sample size for assessment of common genetic variation associated with disease. Additionally, the genotyping was performed several years ago on a platform with limited SNPs (a gene-centric array with 50K SNPs) compared to those now readily available. These limitations mean that PE may share overlapping genetic features with additional traits, but to detect these effects, larger cohorts with higher-coverage genotyping or sequencing will be required. In general, for each trait examined, the power to detect a significant association is dependent on the degree of heritability of the trait, the amount of heritability explained by risk alleles in the literature, and the number of risk alleles assessed on the genotyping array. For example, for hypertension, we demonstrated overlapping genetic architecture between DBP and PE, but not SBP. However, the heritability of SBP and DBP is estimated at ∼20% and 50%, respectively, in European ancestry individuals^82^; thus, we would expect shared genetic architecture of DBP and PE to be demonstrated in a smaller cohort than that required to demonstrate shared genetics of SBP and PE. For specific traits potentially associated with PE (e.g. PCOS, fasting insulin), an assessment could not be performed as there were no proxies on our array for the literature SNPs.

### Interpretation

Prior studies have examined whether genetic predisposition to specific cardiovascular-associated traits (i.e., maternal essential hypertension^83^, dyslipidemia^84^, and CRP^85^) predisposes women to PE. For essential hypertension genetics and PE risk, a similar approach was used and no significant association was found^83^. However, this analysis was done in a smaller cohort (162 PE cases, 108 controls) using risk alleles from a 2011 hypertension GWAS^86^. The genetic instrument included 6 SBP and 6 DBP risk alleles ^83^. In our analysis, an expanded set of risk alleles for SBP (19 SNPs) and DBP (16 SNPs) were used from a subsequent GWAS^35^ in a larger cohort. Thus, we had increased power to detect an association. For dyslipidemia, lower HDL was marginally associated with increased PE risk^84,85^, similar to our observation that risk alleles associated with increased HDL had a suggestive association with decreased PE risk. For the study of CRP (which again used a smaller cohort and fewer SNPs), increased genetic risk for elevated CRP was associated with decreased PE risk^84,85^, which is in contrast to the clinical association of increased CRP with increased PE risk^46^. These findings warrant further interrogation in larger cohorts.

The specific mechanisms by which known clinical risk factors—including hypertension, obesity, and diabetes—contribute to PE development, as well as long-term CVD risk, have not been fully elucidated. Our results suggest that genetic risk factors shared between particular traits (i.e., hypertension, obesity, CAD) and PE may be involved both in the pathogenesis of PE itself and in later CVD development. In published hypertension GWAS, risk loci are enriched for regulatory elements affecting gene expression in vascular endothelial cells and are associated with end organ damage in the heart, cerebral vessels, carotid artery, and the eye^35^. As PE is characterized by diffuse endothelial dysfunction^87^, women with genetic predisposition to altered vascular endothelial cell function are likely to be at high risk for PE. This idea is directly supported by the results of the maternal PE GWAS performed on this same cohort^8^ where the variant, rs9478812 near the gene, *PLEKHG1*, was identified; this locus has been associated with blood pressure in an independent GWAS^88^. In published CAD GWAS, risk loci are highly associated with lipid traits and blood pressure, and pathway analyses highlight lipid metabolism and inflammation as key underlying biological processes^89^, processes also highly associated with PE^24,90–92^. In published obesity GWAS, identified risk loci are highly associated with brain regions important for appetite regulation, learning, emotion, and memory^64^, as well as insulin utilization, energy/lipid metabolism, and adipogenesis in other tissues^64^. Whether these pathways directly contribute to underlying PE pathophysiology or are mediated through BMI-related effects, is an important question for future investigations.

At first glance, the ALP and VTE effects related to PE were surprising, as they were in the opposite directions as anticipated. However, further investigation provides some useful insight. For ALP, increased levels are commonly associated with biliary obstruction, but ALP is also present in bone, intestine, leukocytes, and placenta^93^. Of the 5 SNPs used in the ALP genetic instrument, 2 SNPs drove the association—rs174601 and rs579459. The SNP, rs174601, is associated with decreased *FADS1* and *FADS2* expression in the liver and decreased *FADS1* and increased *FADS2* and *C11orf10* expression in peripheral leukocytes^57^. *FADS1* and *FADS2* are associated with circulating fatty acids in plasma, as well as CAD and ischemic stroke risk^94^. The function of *C11orf10* remains unknown. The SNP, rs579459, is an upstream variant near the *ABO* gene, which encodes for ABO blood group system proteins. C-allele carriers of the rs579459 SNP (corresponding to blood group A) have increased LDL cholesterol^95^ and an increased risk of adverse cardiac outcomes^96^. Thus, the apparent protective nature of SNPs associated with elevated ALP for PE may instead reflect the pleiotropic nature of these SNPs, particularly in relationship to cardiovascular traits.

For VTE, while it is clinically-recognized that women with PE are at significantly increased risk of VTE^42,62^, thrombosis is also a trait highly-associated with recurrent pregnancy loss^97,98^. Given this, women with the highest genetic risk for VTE may have a reduced rate of live births, leading to a decreased expression of all pregnancy-related phenotypes including PE. We hypothesize that this relationship may explain the lower rate of PE seen in women with increased genetic risk for VTE.

Distinct molecular subtypes of PE that go beyond the distinction of early vs. late-onset disease are emerging^99–101^. While much work remains to define these subtypes, our analysis suggests clustering of different trait associations within the broad categories of early vs. late-onset. Specifically, patients with increased genetic risk for elevated DBP are more likely to develop early-onset PE, while patients with increased genetic risk for obesity are more likely to develop term PE. As larger cohorts with more detailed PE phenotyping and genetics emerge, molecular subtyping of PE can be refined further based on both maternal and fetal genetic risk.

## CONCLUSIONS

In conclusion, using curated literature genetic risk loci, we provide the first comprehensive analysis of the overlap of maternal PE genetic architecture with the genetic architecture of other disorders, revealing an overlap of hypertension and obesity genetics with PE. These results implicate underlying genetics as a causal factor for both the pathogenesis of PE itself and the development of later CVD. Expanding understanding of PE heritability will require the establishment of larger maternal and fetal PE consortia with detailed phenotyping and genome-wide genotyping/sequencing. Future analyses should focus not only on the independent effects of the maternal and fetal genomes on disease pathogenesis, but also how the interplay of the two genomes contributes to disease. Such endeavors will allow for more comprehensive delineation of PE heritability and the overlap of PE genetics with the genetic underpinnings of other common diseases.

## Contribution to Authorship

All authors contributed to the conception, design, participant recruitment and data collection in the present study. ACB, KJG, and RS were involved in data analysis. KJG and RS drafted the manuscript. All authors were involved in interpreting the data and critically reviewing the manuscript. All authors gave approval of the final version of the manuscript.

## Details of Ethics Approval

The study was approved under Partners IRB protocol #2011P001053 and #2014P000719, as well as the ethics committees at all participating institutions.

