## Supplemental Tables for "Risk of preeclampsia in patients with genetic predisposition to common medical conditions: a case-control study"

**PRESENT AFFILIATIONS**:

[a] Department of Medicine, Channing Division of Network Medicine, Brigham and Women's Hospital, Boston, MA
[b] Department of Biomedical Informatics, Harvard Medical School, Boston, MA
[c] Department of Anesthesiology, Brigham and Women's Hospital, Boston, MA

### ABBREVIATIONS

OR, odds ratio
SNP, single nucleotide polymorphism

11. Pattaro C, Teumer A, Gorski M, Chu AY, Li M, Mijatovic V, et al. Genetic associations at

53 loci highlight cell types and biological pathways relevant for kidney function. Nat Commun. 2016 Jan 21;7:10023.

### SUPPLEMENTAL TABLES

**Table S1**. Adiponectin: SNPs for genetic risk score calculation

| Literature SNP [1] | Used SNP | $R^2$ | Used SNP: Effect allele | Used SNP: Other allele | β-coefficient for literature SNP |
|---|---|---|---|---|---|
| rs3774261 | rs3774261 | 1.00 | A | G | 0.354 |
| rs3821799 | rs3821799 | 1.00 | C | T | 0.352 |
| rs17300539 | rs822387 | 0.83 | C | T | 0.330 |

**Table S2**. Alkaline phosphatase (ALP): SNPs for genetic risk score calculation*

| Literature SNP [2] | Used SNP | $R^2$ | Used SNP: Effect allele | Used SNP: Other allele | β-coefficient for literature SNP |
|---|---|---|---|---|---|
| rs16856332 | rs10199694 | 1.00 | A | G | 3.9 |
| rs174601 | rs174601 | 1.00 | C | T | 1.7 |
| rs579459 | rs651007 | 1.00 | T | C | 8.8 |
| rs2954021 | rs2980875 | 0.78 | G | A | 1.4 |
| rs7267979 | rs6083776 | 0.59 | G | A | 1.5 |

*The following literature SNPs were not analyzed since there were no proxies available: rs10819937, rs1883415, rs1976403, rs2236653, rs281377, rs314253, rs6984305, rs7186908, rs7923609.

**Table S3**. Birthweight: SNPs for genetic risk score calculation*

| Literature SNP [3] | Used SNP | $R^2$ | Used SNP: Effect allele | Used SNP: Other allele | β-coefficient for literature SNP |
|---|---|---|---|---|---|
| rs1351394 | rs1042725 | 0.84 | T | C | 0.023 |
| rs35261542 | rs9368222 | 1.00 | C | A | 0.044 |
| rs7076938 | rs7076938 | 1.00 | T | C | 0.036 |
| rs700059 | rs2488597 | 0.80 | G | A | 0.033 |
| rs12823128 | rs7309939 | 0.77 | T | C | 0.021 |
| rs10830963 | rs10830963 | 1.00 | G | C | 0.023 |

*The following literature SNPs were not analyzed since there were no proxies available: rs13322435, rs1171920, rs925098, rs854037, rs1374204, rs138715366, rs1101081, rs28510415, rs113086489, rs61862780, rs6537307, rs3753639, rs62240962, rs62466330, rs13266210, rs1415701, rs7575873, rs28530618, rs6016377, rs10935733, rs72851023, rs11765649, rs2473248, rs7402982, rs11096402, rs7729301, rs144843919, rs72480273, rs1819436, rs74233809, rs12543725, rs1011939, rs12942207, rs798489, rs7847628, rs2421016, rs6040076, rs2324499, rs7964361, rs12906125, rs6959887, rs7742369, rs9379832, rs2242116, rs2229742, rs2854355, rs134594, rs11055034, rs10402712, rs61154119, rs2150052, rs61830764, rs6989280, rs139975827.

**Table S4**. Body mass index (BMI): SNPs for genetic risk score calculation*

| Literature SNP [4] | Used SNP | $R^2$ | Used SNP: Effect allele | Used SNP: Other allele | β-coefficient for literature SNP |
|---|---|---|---|---|---|
| rs1558902 | rs1421085 | 1.00 | C | T | 0.079 |
| rs12566985 | rs7553158 | 1.00 | G | A | 0.027 |
| rs3888190 | rs4788099 | 1.00 | G | A | 0.028 |
| rs17001654 | rs17001561 | 1.00 | A | G | 0.032 |
| rs12705981 | rs12705973 | 1.00 | G | A | 0.021 |
| rs2075650 | rs2075650 | 1.00 | A | G | 0.031 |
| rs2246012 | rs2246012 | 1.00 | C | T | 0.022 |
| rs1808579 | rs2236707 | 0.97 | C | T | 0.022 |
| rs9880211 | rs9844666 | 0.95 | A | G | 0.023 |
| rs11030104 | rs10767664 | 0.91 | A | T | 0.038 |
| rs12016871 | rs7988412 | 0.78 | T | C | 0.031 |
| rs16951275 | rs6494696 | 0.72 | G | C | 0.030 |
| rs2287019 | rs10423928 | 0.67 | T | A | 0.032 |
| rs3817334 | rs11570094 | 0.66 | A | C | 0.027 |
| rs2112347 | rs6453133 | 0.57 | A | G | 0.030 |

*The following literature SNPs were not analyzed since there were no proxies available: rs11672550, rs13078960, rs12450239, rs17024393, rs1928295, rs11917972, rs7550169, rs29941, rs9856151, rs1830074, rs11247009, rs268073, rs2579103, rs10938353, rs6062788, rs12885454, rs1464321, rs4566392, rs38313, rs2650492, rs4984406, rs1979755, rs657452, rs9867325.

**Table S5**. Coronary artery disease (CAD): SNPs for genetic risk score calculation*

| Literature SNP [5,6] | Used SNP | $R^2$ | Used SNP: Effect allele | Used SNP: Other allele | OR for literature SNP |
|---|---|---|---|---|---|
| rs11206510 | rs11206510 | 1.00 | T | C | 1.08 |
| rs11556924 | rs11556924 | 1.00 | C | T | 1.08 |
| rs17514846 | rs17514846 | 1.00 | A | C | 1.05 |
| rs2519093 | rs651007 | 0.96 | T | C | 1.08 |
| rs2681472 | rs11105354 | 1.00 | G | A | 1.08 |
| rs2891168 | rs2891168 | 1.00 | G | A | 1.21 |
| rs2954029 | rs10808546 | 1.00 | C | T | 1.04 |
| rs3184504 | rs3184504 | 1.00 | T | C | 1.07 |
| rs7528419 | rs7528419 | 1.00 | A | G | 1.12 |
| rs2244608 | rs2244608 | 1.00 | G | A | 1.06 |

*The following literature SNPs were not analyzed since there were no proxies available: rs10139550, rs10840293, rs11191416, rs11838776, rs12202017, rs1412444, rs16986953, rs17087335, rs17678683, rs180803, rs1870634, rs2107595, rs2128739, rs216172, rs2487928, rs28451064, rs3918226, rs4252185, rs4420638, rs4468572, rs4593108, rs515135, rs55730499, rs56062135, rs56289821, rs56336142, rs6544713, rs663129, rs6689306, rs67180937, rs6725887, rs7212798, rs72689147, rs7568458, rs8042271, rs9349379, rs9818870, rs9970807, rs11810571, rs7623687, rs142695226, rs433903, rs10857147, rs11723436, rs35879803, rs1351525, rs11170820, rs7500448, rs8108632.

**Table S6**. Total cholesterol: SNPs for genetic risk score calculation*

| **Literature SNP** [7,8] | **Used SNP** | $R^2$ | **Used SNP: Effect allele** | **Used SNP: Other allele** | **β-coefficient for literature SNP** |
|---|---|---|---|---|---|
| rs10401969 | rs3794991 | 1.00 | T | C | -0.137 |
| rs12027135 | rs873308 | 1.00 | G | A | -0.027 |
| rs1260326 | rs1260326 | 1.00 | T | C | 0.051 |
| rs12916 | rs12916 | 1.00 | C | T | 0.068 |
| rs1367117 | rs1367117 | 1.00 | A | G | 0.100 |
| rs1800961 | rs1800961 | 1.00 | T | C | -0.106 |
| rs1883025 | rs1883025 | 1.00 | T | C | -0.067 |
| rs1961456 | rs1961456 | 1.00 | G | A | 0.032 |
| rs2000999 | rs2000999 | 1.00 | A | G | 0.062 |
| rs2479409 | rs2479409 | 1.00 | G | A | 0.054 |
| rs3764261 | rs3764261 | 1.00 | A | C | 0.050 |
| rs4299376 | rs4299376 | 1.00 | G | T | 0.079 |
| rs629301 | rs646776 | 1.00 | C | T | -0.134 |
| rs651007 | rs651007 | 1.00 | T | C | 0.069 |
| rs6511720 | rs6511720 | 1.00 | T | G | -0.185 |
| rs174550 | rs1535 | 0.97 | A | G | -0.048 |
| rs3850634 | rs1748197 | 0.97 | A | G | -0.075 |
| rs2290159 | rs9817675 | 0.80 | T | C | -0.037 |
| rs964184 | rs2075290 | 0.63 | T | C | -0.121 |
| rs1532085 | rs4775041 | 0.57 | C | G | 0.054 |

*The following literature SNPs were not analyzed since there were no proxies available: rs10832963, rs11065987, rs11136341, rs11220463, rs1169288 , rs117087731, rs12983728, rs1564348, rs1800562, rs2072183 , rs2081687 , rs2126259, rs2255141, rs2277862, rs2285942, rs2807834, rs2814982 , rs2902940, rs2954022, rs3757354, rs4297946, rs4420638 , rs492602, rs514230, rs581080, rs6759321, rs6882076, rs7206971, rs7239867, rs7515577, rs7941030, rs9488822.

**Table S7**. C-reactive protein (CRP): SNPs for genetic risk score calculation*

| **Literature SNP** [9] | **Used SNP** | $R^2$ | **Used SNP: Effect allele** | **Used SNP: Other allele** | **β-coefficient for literature SNP** |
|---|---|---|---|---|---|
| rs12239046 | rs12239046 | 1.00 | C | T | 0.048 |
| rs1260326 | rs1260326 | 1.00 | T | C | 0.089 |
| rs1800961 | rs1800961 | 1.00 | C | T | 0.120 |
| rs2794520 | rs1205 | 1.00 | C | T | 0.193 |
| rs4129267 | rs4129267 | 1.00 | C | T | 0.094 |
| rs4420065 | rs1805096 | 1.00 | G | A | 0.111 |
| rs1183910 | rs2244608 | 0.96 | A | G | 0.152 |
| rs13233571 | rs13232120 | 0.93 | A | T | 0.054 |

| | | | | | |
|---|---|---|---|---|---|
| rs6734238 | rs4251961 | 0.61 | C | T | 0.047 |

*The following literature SNPs were not analyzed since there were no proxies available: rs10521222, rs10745954, rs12037222, rs2836878, rs340029, rs4420638, rs4903031, rs9987289.

**Table S8**. Diastolic blood pressure (DBP): SNPs for genetic risk score calculation*

| Literature SNP [10] | Used SNP | $R^2$ | Used SNP: Effect allele | Used SNP: Other allele | β-coefficient for literature SNP |
|---|---|---|---|---|---|
| rs11105354 | rs11105354 | 1.00 | A | G | 0.459 |
| rs11556924 | rs11556924 | 1.00 | T | C | -0.214 |
| rs13306560 | rs13306560 | 1.00 | T | C | 0.290 |
| rs1450271 | rs7944706 | 0.76 | A | G | 0.199 |
| rs1620668 | rs1620668 | 1.00 | A | G | -0.285 |
| rs16823124 | rs16823124 | 1.00 | A | G | 0.265 |
| rs17037390 | rs17037390 | 1.00 | A | G | -0.499 |
| rs17367504 | rs17367504 | 1.00 | A | G | 0.390 |
| rs2493134 | rs2493134 | 1.00 | T | C | -0.275 |
| rs3184504 | rs3184504 | 1.00 | T | C | 0.362 |
| rs4494250 | rs10786172 | 0.93 | A | G | 0.150 |
| rs4746172 | rs3793921 | 1.00 | T | C | 0.076 |
| rs4846049 | rs4846049 | 1.00 | T | G | -0.196 |
| rs592373 | rs592373 | 1.00 | A | G | 0.282 |
| rs6271 | rs6271 | 1.00 | T | C | -0.465 |
| rs936226 | rs7085 | 0.85 | T | C | -0.363 |

*The following literature SNPs were not analyzed since there were no proxies available: rs10077885, rs10850411, rs10943605, rs10995311, rs10995311, rs11014166, rs110419, rs11128722, rs1126464, rs1126464, rs1156725, rs11953630, rs12243859, rs12521868, rs12627651, rs12656497, rs12940887, rs12958173, rs13082711, rs13107325, rs13139571, rs1327235, rs1361831, rs1371182, rs1378942, rs1401454, rs1458038, rs167479, rs17030613, rs17080093, rs17428471, rs17638167, rs1799945, rs1975487, rs2187668, rs2272007, rs2291435, rs2384550, rs2521501, rs2586886, rs2891546, rs2898290, rs2932538, rs2969070, rs35444, rs3735533, rs3752728, rs381815, rs4245739, rs4247374, rs4590817, rs4660293, rs6026748, rs6095241, rs633185, rs6442101, rs6495122, rs6779380, rs6825911, rs687621, rs689134, rs6969780, rs7076398, rs7103648, rs711737, rs7213273, rs7302981, rs7302981, rs740746, rs7515635, rs751984, rs76452347, rs772178, rs805303, rs8068318, rs8068318, rs880315, rs891511, rs900145, rs918466, rs926552, rs932764, rs943037, rs9687065.

**Table S9**. Estimated glomerular filtration rate from serum creatinine (eGFRCr): SNPs for genetic risk score calculation*

| Literature SNP [11] | Used SNP | $R^2$ | Used SNP: Effect allele | Used SNP: Other allele | β-coefficient for literature SNP |
|---|---|---|---|---|---|
| rs8091180 | rs8091180 | 1.00 | A | G | -0.006 |
| rs12136063 | rs11142 | 0.89 | A | G | 0.0045 |
| rs4667594 | rs10490132 | 0.91 | C | A | -0.0044 |
| rs1260326 | rs1260326 | 1.00 | T | C | 0.0068 |

| | | | | | |
|---|---|---|---|---|---|
| rs6546838 | rs6546838 | 1.00 | A | G | -0.0093 |
| rs9472135 | rs9369425 | 0.77 | A | G | -0.008 |
| rs316009 | rs316009 | 1.00 | T | C | 0.0131 |
| rs7805747 | rs10480300 | 1.00 | T | C | -0.013 |
| rs3758086 | rs3758086 | 1.00 | A | G | -0.0071 |

*The following literature SNPs were not analyzed since there were no proxies available: rs3850625, rs2712184, rs9682041, rs10513801, rs10994860, rs163160, rs164748, rs2802729, rs6795744, rs228611, rs7759001, rs10277115, rs3750082, rs6459680, rs4014195, rs10491967, rs7956634, rs1106766, rs11666497, rs6088580, rs17216707, rs1800615, rs267734, rs807601, rs7422339, rs2861422, rs17319721, rs11959928, rs6420094, rs848490, rs4744712, rs1044261, rs963837, rs10774021, rs716877, rs476633, rs2467853, rs491567, rs1394125, rs13329952, rs2453580, rs9916302, rs11657044, rs12460876.

**Table S10**. Fibrinogen: SNPs for genetic risk score calculation*

| Literature SNP [12] | Used SNP | $R^2$ | Used SNP: Effect allele | Used SNP: Other allele | β-coefficient for literature SNP |
|---|---|---|---|---|---|
| rs3138493 | rs1571536 | 0.91 | T | C | -0.005 |
| rs73058052 | rs2304204 | 0.56 | T | C | 0.007 |
| rs10157379 | rs12239046 | 0.93 | C | T | -0.010 |
| rs59104589 | rs15129 | 0.52 | T | C | -0.008 |
| rs1800961 | rs1800961 | 1.00 | T | C | -0.017 |

*The following literature SNPs were not analyzed since there were no proxies available: rs7588285, rs62246343, rs1976714, rs2710804, rs7012814, rs2250644, rs2420915, rs7934094, rs7310615, rs56702977, rs1035560, rs7224737, rs1892534, rs61812598, rs1558643, rs6734238, rs715, rs9840812, rs59950280, rs7439150, rs2057655, rs71520386, rs11780978, rs7916868, rs11230201, rs2731439, rs367677, rs12913259, rs11859517, rs9808651, rs75347843.

**Table S11**. Fasting glucose: SNPs for genetic risk score calculation*

| Literature SNP [1] | Used SNP | $R^2$ | Used SNP: Effect allele | Used SNP: Other allele | β-coefficient for literature SNP |
|---|---|---|---|---|---|
| rs10830963 | rs10830963 | 1.00 | G | C | 0.067 |
| rs4607517 | rs1799884 | 1.00 | T | C | 0.062 |
| rs11920090 | rs10513685 | 0.93 | G | A | 0.020 |
| rs7944584 | rs11039149 | 0.84 | A | G | 0.021 |
| rs560887 | rs569805 | 0.73 | T | A | 0.075 |

*The following literature SNPs were not analyzed since there were no proxies available: rs10885122, rs11071657, rs11605924, rs11708067, rs13266634, rs2191349, rs340874, rs7034200.

**Table S12**. Gamma-glutamyl transferase (GTT): SNPs for genetic risk score calculation*

| Literature SNP [2] | Used SNP | $R^2$ | Used SNP: Effect allele | Used SNP: Other allele | β-coefficient for literature SNP |
|---|---|---|---|---|---|
| rs10513686 | rs11711437 | 1.00 | C | G | 4.9 |

| | | | | | |
|---|---|---|---|---|---|
| rs1260326 | rs1260326 | 1.00 | C | T | 3.2 |
| rs17145750 | rs17145750 | 1.00 | T | C | 4.5 |
| rs12145922 | rs786918 | 0.82 | A | G | 2.8 |
| rs2739330 | rs1006771 | 0.80 | T | G | 3.7 |
| rs7310409 | rs2244608 | 0.62 | G | A | 6.8 |

*The following literature SNPs were not analyzed since there were no proxies available: rs1076540, rs10908458, rs12968116, rs13030978, rs1335645, rs1497406, rs2073398, rs2140773, rs339969, rs4074793, rs4503880, rs4547811, rs4581712, rs516246, rs6888304, rs754466, rs8038465, rs9296736, rs944002, rs9913711.

**Table S13**. High-density lipoprotein (HDL): SNPs for genetic risk score calculation*

| Literature SNP [7,8] | Used SNP | $R^2$ | Used SNP: Effect allele | Used SNP: Other allele | β-coefficient for literature SNP |
|---|---|---|---|---|---|
| rs1121980 | rs1121980 | 1.00 | A | G | -0.020 |
| rs12678919 | rs10503669 | 1.00 | A | C | 0.155 |
| rs13326165 | rs13326165 | 1.00 | A | G | 0.029 |
| rs17145738 | rs13232120 | 1.00 | T | A | 0.041 |
| rs174601 | rs174601 | 1.00 | T | C | -0.039 |
| rs1800961 | rs1800961 | 1.00 | T | C | -0.127 |
| rs1883025 | rs1883025 | 1.00 | T | C | -0.070 |
| rs3136441 | rs2070850 | 1.00 | T | C | 0.054 |
| rs3764261 | rs3764261 | 1.00 | A | C | 0.241 |
| rs4846914 | rs4846914 | 1.00 | G | A | -0.048 |
| rs6065906 | rs6073952 | 1.00 | A | G | -0.059 |
| rs6805251 | rs6782799 | 1.00 | T | C | 0.020 |
| rs881844 | rs881844 | 1.00 | G | C | -0.032 |
| rs7255436 | rs2278236 | 0.97 | G | A | -0.032 |
| rs838880 | rs838878 | 0.96 | A | G | 0.048 |
| rs7241918 | rs2156552 | 0.95 | A | T | -0.090 |
| rs7134594 | rs7298565 | 0.94 | A | G | -0.035 |
| rs2013208 | rs1062633 | 0.90 | T | C | 0.025 |
| rs386000 | rs103294 | 0.89 | T | C | 0.048 |
| rs2652834 | rs11071721 | 0.85 | T | C | -0.028 |
| rs16942887 | rs20549 | 0.79 | G | A | 0.083 |
| rs2606736 | rs347591 | 0.79 | G | T | 0.025 |
| rs3741414 | rs11172134 | 0.72 | A | T | 0.028 |
| rs970548 | rs1487562 | 0.69 | T | C | 0.026 |
| rs12145743 | rs928391 | 0.68 | C | T | 0.020 |
| rs964184 | rs2075290 | 0.63 | T | C | 0.106 |
| rs1532085 | rs4775041 | 0.57 | C | G | 0.107 |
| rs12801636 | rs3741378 | 0.56 | T | C | 0.024 |

*The following literature SNPs were not analyzed since there were no proxies available: rs10019888, rs1047891, rs10808546, rs11246602, rs12328675, rs12748152, rs12967135, rs13107325, rs1515100, rs1689800, rs17173637, rs17695224, rs181362, rs1936800, rs2290547, rs2293889, rs2602836, rs2923084, rs2925979, rs3822072, rs4082919, rs4142995, rs4148008, rs4420638, rs4650994, rs4660293, rs4731702, rs4759375, rs4765127, rs4917014, rs4983559, rs499974, rs605066, rs643531, rs6450176, rs702485, rs7115089, rs7134375, rs7188861, rs731839, rs737337, rs78058190, rs998584, rs9987289.

**Table S14**. Ischemic stroke: SNPs for genetic risk score calculation*

| Literature SNP [13] | Used SNP | $R^2$ | Used SNP: Effect allele | Used SNP: Other allele | OR for literature SNP |
|---|---|---|---|---|---|
| rs6843082 | rs4611994 | 0.53 | C | T | 1.36 |
| rs2383207 | rs944797 | 1.00 | C | T | 1.15 |
| rs505922 | rs657152 | 1.00 | A | C | 1.13 |
| rs660599 | rs17368659 | 0.55 | T | G | 1.18 |
| rs10744777 | rs10744777 | 1.00 | T | C | 1.10 |

*The following literature SNPs were not analyzed since there were no proxies available: rs556621, rs2107595, rs879324.

**Table S15**. Low-density lipoprotein (LDL): SNPs for genetic risk score calculation*

| Literature SNP [7,8] | Used SNP | $R^2$ | Used SNP: Effect allele | Used SNP: Other allele | β-coefficient for literature SNP |
|---|---|---|---|---|---|
| rs10401969 | rs3794991 | 1.00 | T | C | -0.118 |
| rs12027135 | rs873308 | 1.00 | G | A | -0.030 |
| rs12916 | rs12916 | 1.00 | C | T | 0.073 |
| rs1367117 | rs1367117 | 1.00 | A | G | 0.119 |
| rs174583 | rs174576 | 1.00 | A | C | -0.051 |
| rs2000999 | rs2000999 | 1.00 | A | G | 0.065 |
| rs2479409 | rs2479409 | 1.00 | G | A | 0.064 |
| rs4299376 | rs4299376 | 1.00 | G | T | 0.081 |
| rs629301 | rs646776 | 1.00 | C | T | -0.167 |
| rs649129 | rs651007 | 1.00 | T | C | 0.077 |
| rs3850634 | rs1748197 | 0.97 | A | G | -0.049 |
| rs964184 | rs2075290 | 0.63 | T | C | -0.086 |

*The following literature SNPs were not analyzed since there were no proxies available: rs11065987, rs11136341, rs11153594, rs11220462, rs1129555, rs1169288, rs117492019, rs12670798, rs1564348, rs1800562, rs182616603, rs2081687, rs2126259, rs217386, rs247616, rs2807834, rs2902941, rs2954022, rs3177928, rs3757354, rs4420638, rs514230, rs6511720, rs6882076, rs7206971, rs79588679, rs8017377, rs909802.

**Table S16**. Systolic blood pressure (SBP): SNPs for genetic risk score calculation*

| Literature SNP [10] | Used SNP | $R^2$ | Used SNP: Effect allele | Used SNP: Other allele | β-coefficient for literature SNP |
|---|---|---|---|---|---|
| rs10224002 | rs10224002 | 1.00 | A | G | -0.465 |
| rs11105354 | rs11105354 | 1.00 | A | G | 0.909 |
| rs11229457 | rs11229519 | 0.74 | T | C | -0.310 |

| | | | | | |
|---|---|---|---|---|---|
| rs12247028 | rs7080350 | 0.56 | A | G | -0.364 |
| rs1450271 | rs7944706 | 0.76 | A | G | 0.413 |
| rs1620668 | rs1620668 | 1.00 | A | G | -0.535 |
| rs17367504 | rs17367504 | 1.00 | A | G | 0.716 |
| rs217727 | rs217727 | 1.00 | A | G | 0.591 |
| rs2493134 | rs2493134 | 1.00 | T | C | -0.431 |
| rs3184504 | rs3184504 | 1.00 | T | C | 0.498 |
| rs347591 | rs347591 | 1.00 | T | G | 0.244 |
| rs3741378 | rs3741378 | 1.00 | T | C | -0.486 |
| rs5219 | rs5215 | 0.94 | T | C | 0.320 |
| rs592373 | rs592373 | 1.00 | A | G | 0.484 |
| rs6271 | rs6271 | 1.00 | T | C | -0.591 |
| rs7129220 | rs4403799 | 0.92 | A | G | -1.107 |
| rs7297416 | rs7297416 | 1.00 | A | C | 0.090 |
| rs757081 | rs5215 | 0.65 | T | C | -0.085 |
| rs936226 | rs7085 | 0.85 | T | C | -0.549 |

*The following literature SNPs were not analyzed since there were no proxies available:
rs10077885, rs10760117, rs111245230, rs11128722, rs11556924a, rs1156725, rs11639856, rs1173766, rs11953630, rs12243859, rs12279202, rs12627651, rs12656497, rs12705390, rs12940887, rs12940887, rs12946454, rs12958173, rs13107325, rs1327235, rs13359291, rs1361831, rs1371182, rs1378942, rs1458038, rs16849225, rs16877320, rs17010957, rs17037390, rs17080093, rs17365948, rs17428471, rs17608766, rs17638167, rs1799945, rs1813353, rs1975487, rs2014912, rs2270860, rs2291435, rs2493292, rs2521501, rs2586886, rs2594992, rs2891546, rs2898290, rs2932538, rs2969070, rs35479618, rs35529250, rs3735533, rs3752728, rs381815, rs4245739, rs4247374, rs4387287, rs4590817, rs4691707, rs4823006, rs6026748, rs61760904, rs633185, rs6442101, rs6722745, rs6779380, rs689134, rs6919440, rs7076398, rs7103648, rs711737, rs7213273, rs740746, rs7515635, rs751984, rs805303, rs880315, rs932764, rs9349379, rs943037.

**Table S17**. Systemic lupus erythematosus (SLE): SNPs for genetic risk score calculation*

| **Literature SNP** [14] | **Used SNP** | **$R^2$** | **Used SNP: Effect allele** | **Used SNP: Other allele** | **OR for literature SNP** |
|---|---|---|---|---|---|
| rs10036748 | rs7708392 | 1.00 | C | G | 1.33 |
| rs10488631 | rs10488631 | 1.00 | C | T | 1.84 |
| rs2286672 | rs2286672 | 1.00 | T | C | 1.24 |
| rs2476601 | rs2476601 | 1.00 | A | G | 1.37 |
| rs3794060 | rs7928249 | 1.00 | A | G | 1.12 |
| rs9652601 | rs2041670 | 0.96 | G | A | 1.13 |
| rs10774625 | rs3184504 | 0.93 | T | C | 1.17 |
| rs6932056 | rs2230926 | 0.79 | G | T | 1.70 |
| rs34572943 | rs13338069 | 0.69 | G | A | 1.69 |

*The following literature SNPs were not analyzed since there were no proxies available:
rs10028805, rs1059312, rs11644034, rs11889341, rs1270942, rs12802200, rs1734787, rs17849501, rs1801274, rs2111485, rs2289583, rs2304256, rs2431697, rs2663052, rs2732549,

rs2736340, rs2941509, rs3024505, rs3768792, rs4902562, rs4917014, rs4948496, rs564799, rs6568431, rs6740462, rs704840, rs7444, rs7726414, rs7941765, rs849142, rs887369, rs9311676, rs9462027, rs9782955.

**Table S18**. Type 2 diabetes mellitus (T2DM): SNPs for genetic risk score calculation*

| Literature SNP [15] | Used SNP | $R^2$ | Used SNP: Effect allele | Used SNP: Other allele | OR for literature SNP |
|---|---|---|---|---|---|
| rs6757251 | rs7578597 | 1.00 | C | T | 1.14 |
| rs4402960 | rs4402960 | 1.00 | T | G | 1.15 |
| rs3821943 | rs3821943 | 1.00 | T | C | 1.10 |
| rs1635852 | rs864745 | 0.97 | T | C | 1.10 |
| rs878521 | rs2908282 | 0.55 | A | G | 1.05 |
| rs3802177 | rs3802177 | 1.00 | G | A | 1.12 |
| rs10757282 | rs10757282 | 1.00 | C | T | 1.04 |
| rs635634 | rs651007 | 1.00 | T | C | 1.08 |
| rs7903146 | rs7903146 | 1.00 | T | C | 1.34 |
| rs233449 | rs233449 | 1.00 | G | A | 1.09 |
| rs5219 | rs5215 | 0.94 | T | C | 1.07 |
| rs76550717 | rs613937 | 0.79 | A | G | 1.10 |
| rs10830963 | rs10830963 | 1.00 | G | C | 1.08 |
| rs6581998 | rs4760921 | 0.70 | T | A | 1.06 |
| rs56348580 | rs12427353 | 0.66 | G | C | 1.08 |
| rs78761021 | rs7219033 | 0.87 | G | A | 1.07 |
| rs757209 | rs4239217 | 0.63 | G | A | 1.09 |
| rs12454712 | rs12454712 | 1.00 | T | C | 1.05 |
| rs58489806 | rs3794991 | 0.64 | T | C | 1.09 |
| rs12625671 | rs4810426 | 0.87 | C | T | 1.09 |
| rs1800961 | rs1800961 | 1.00 | T | C | 1.17 |

*The following literature SNPs were not analyzed since there were no proxies available: rs3768321, rs12031920, rs406767, rs67156297, rs340874, rs145819220, rs9309245, rs1116357, rs10193447, rs6723108, rs7560163, rs1563575, rs28584669, rs2972156, rs1861612, rs11712037, rs35352848, rs79819696, rs7428936, rs11708067, rs9820223, rs6777684, rs1531583, rs7660590, rs60780116, rs11747901, rs9687833, rs173964, rs6453287, rs78408340, rs74944275, rs6923241, rs7451008, rs115321690, rs2244020, rs9271774, rs143308245, rs139514607, rs11759026, rs6918311, rs10276674, rs10238625, rs10229583, rs73455744, rs791595, rs10954284, rs1182436, rs516946, rs4734285, rs11786613, rs10758593, rs186838848, rs10965248, rs10965223, rs1575972, rs13301067, rs9410573, rs11787792, rs11257659, rs10998572, rs810517, rs11187140, rs10886471, rs2292626, rs2334499, rs11564732, rs7107784, rs756852, rs231360, rs2237897, rs191294997, rs441613, rs1061810, rs111669836, rs11063018, rs188827514, rs4238013, rs7953190, rs147538848, rs2258238, rs2851437, rs9552911, rs7330796, rs11616380, rs10146997, rs7403531, rs67839313, rs4774420, rs952471, rs62006309, rs62023387, rs12595616, rs1558902, rs8056814, rs2925979, rs9911305, rs7224685, rs13342692, rs79349575, rs7234111, rs1942880, rs79851087, rs139990642, rs429358, rs55864746, rs2023681, rs5945326.

**Table S19**. Triglycerides: SNPs for genetic risk score calculation*

| Literature SNP [7,8] | Used SNP | $R^2$ | Used SNP: Effect allele | Used SNP: Other allele | β-coefficient for literature SNP |
|---|---|---|---|---|---|
| rs10401969 | rs3794991 | 1.00 | T | C | -0.121 |
| rs1260326 | rs1260326 | 1.00 | T | C | 0.115 |
| rs12678919 | rs10503669 | 1.00 | A | C | -0.170 |
| rs1321257 | rs2281719 | 1.00 | C | T | 0.040 |
| rs174546 | rs1535 | 1.00 | G | A | 0.045 |
| rs2131925 | rs1748197 | 1.00 | A | G | -0.066 |
| rs2954029 | rs10808546 | 1.00 | T | C | -0.076 |
| rs4810479 | rs4810479 | 1.00 | C | T | 0.053 |
| rs7248104 | rs7508679 | 1.00 | T | C | -0.022 |
| rs2068888 | rs4418728 | 0.97 | T | G | -0.024 |
| rs442177 | rs3775214 | 0.93 | G | A | -0.031 |
| rs11613352 | rs11172134 | 0.77 | A | T | -0.028 |
| rs5756931 | rs4820314 | 0.66 | A | G | -0.020 |
| rs964184 | rs2075290 | 0.63 | T | C | -0.234 |
| rs11649653 | rs749767 | 0.62 | G | A | -0.027 |

*The following literature SNPs were not analyzed since there were no proxies available: rs10761731, rs11776767, rs12310367, rs13238203, rs1495743, rs1553318, rs1832007, rs193042029, rs2255811, rs2412710, rs2540948, rs261342, rs2929282, rs2943645, rs3198697, rs340839, rs38855, rs645040, rs6831256, rs7205804, rs731839, rs7811265, rs8077889, rs9686661, rs998584.

**Table S20**. Uric acid: SNPs for genetic risk score calculation*

| Literature SNP [16] | Used SNP | $R^2$ | Used SNP: Effect allele | Used SNP: Other allele | β-coefficient for literature SNP |
|---|---|---|---|---|---|
| rs1260326 | rs1260326 | 1.00 | T | C | 0.074 |
| rs2231142 | rs2231142 | 1.00 | T | G | 0.217 |
| rs1178977 | rs17145713 | 1.00 | C | T | 0.047 |
| rs10480300 | rs10480300 | 1.00 | T | C | 0.035 |
| rs6598541 | rs4966021 | 0.96 | C | T | 0.043 |
| rs17786744 | rs3758086 | 0.90 | G | A | -0.029 |
| rs653178 | rs3184504 | 0.87 | C | T | -0.035 |
| rs7193778 | rs33063 | 0.85 | G | A | -0.046 |
| rs729761 | rs9369425 | 0.75 | G | A | -0.047 |
| rs3741414 | rs11172134 | 0.72 | A | T | -0.072 |
| rs11264341 | rs4072037 | 0.55 | C | T | -0.050 |

*The following literature SNPs were not analyzed since there were no proxies available: rs1471633, rs12498742, rs675209, rs1165151, rs1171614, rs2078267, rs478607, rs17050272, rs2307394, rs6770152, rs17632159, rs2941484, rs10821905, rs642803, rs1394125, rs7188445, rs7224610, rs2079742, rs164009.

**Table S21**. Venous thromboembolism (VTE): SNPs for genetic risk score calculation*

| Literature SNP [17] | Used SNP | $R^2$ | Used SNP: Effect allele | Used SNP: Other allele | OR for literature SNP |
|---|---|---|---|---|---|
| rs6025 | rs6025 | 1.00 | T | C | 3.25 |
| rs4524 | rs4524 | 1.00 | T | C | 1.20 |
| rs2066865 | rs2066861 | 1.00 | T | C | 1.24 |
| rs529565 | rs657152 | 1.00 | A | C | 1.55 |
| rs4253417 | rs3756009 | 0.80 | G | A | 1.27 |

*The following literature SNPs were not analyzed since there were no proxies available: rs1799963, rs6087685, rs4602861, rs78707713, rs2288904.
